# Haplotype-aware long-read error correction

**DOI:** 10.1101/2025.06.23.661108

**Authors:** Parvesh Barak, Daniel Gibney, Chirag Jain

## Abstract

Error correction of long reads is an important initial step in genome assembly workflows. For organisms with ploidy greater than one, it is important to preserve haplotype-specific variation during read correction. This challenge has driven the development of several haplotype-aware correction methods. However, existing methods are based on either ad-hoc heuristics or deep learning approaches. In this paper, we introduce a rigorous formulation for this problem. Our approach builds on the minimum error correction framework used in reference-based haplotype phasing. We prove that the proposed formulation for error correction of reads in *de novo* context, i.e., without using a reference genome, is NP-hard. To make our exact algorithm scale to large datasets, we introduce practical heuristics. Experiments using PacBio HiFi sequencing datasets from human and plant genomes show that our approach achieves accuracy comparable to state-of-the-art methods. The software is freely available at https://github.com/at-cg/HALE.

## 1 Introduction

Recent improvements in long-read sequencing technologies in terms of read length and accuracy have led to a paradigm shift in the quality of genome assemblies [14]. Currently, Oxford Nanopore Technology (ONT) long reads have error rates between 1% and 5%, while Pacific Biosciences (PacBio) reads have lower error rates below 0.5% [8]. Error correction is important for distinguishing haplotypes and repeats during genome assembly [20]. The key challenge is that the sequencing error rates in long reads often exceed the variance between repetitive genomic regions that we want to separate.

It is possible to correct sequencing errors by leveraging redundancy in sequencing data because each base in the genome is sampled by multiple reads during whole-genome sequencing [23]. One simple approach is to consider all-vs-all overlaps among reads, and subsequently correct each read by doing a majority vote among the bases of reads that overlap it. Unfortunately, this direct strategy leads to a loss of haplotype-specific and repeat-specific variation. As an example, if two genomic regions of a haploid genome differ at only a few positions, then their corresponding reads may overlap with each other. The majority vote strategy in such cases would eliminate the true biological differences. This issue becomes even more pronounced in diploid and polyploid genomes. In humans, the two haplotypes are approximately 99.9% identical. As a result, reads from different haplotypes overlap frequently. Complicating matters even further, both sequencing errors and biological variations are unevenly distributed across the genome. Certain regions of a genome, such as homopolymers, have high error rates [11].

Long-read assemblers that support diploid genome assembly [1, 2, 3, 13] incorporate haplotype-aware error correction as a mandatory step in their pipelines. More recently, dedicated tools for error correction [10, 18] have also been developed. Most algorithms begin by computing all-vs-all overlaps between reads. For instance, given a target read to be corrected, Hifiasm [2] considers the alignments of overlapping reads to that target read. It uses heuristics to identify heterozygous loci in the target read. Subsequently, it discards those overlapping reads that do not match the target read at all the heterozygous loci. Herro [18] is a dedicated error correction tool that uses a supervised learning model composed of convolutional blocks and a transformer encoder. PECAT [13] uses a heuristic scoring strategy based on partial-order alignment graph [6] for each read to be corrected. DeChat [10] also adopts a heuristic-driven method, performing an initial pre-correction using a de Bruijn graph, followed by a variant-aware multiple sequence alignment strategy adapted from Hifiasm.

In this work, we introduce a rigorous mathematical formulation of the haplotype-aware error correction problem. For each target read, we define a combinatorial optimization problem aimed at selecting reads that originate from the same haplotype and genomic region. An accurate selection of such reads enables a direct consensus-based correction using these reads. We prove that this optimization problem is NP-hard via a reduction from the Max-Cut problem. Despite its hardness, we show that it can be practically solved for human genome datasets using a combination of simple heuristics and brute-force search. We implemented our approach in a tool named HALE (**H**aplotype-**a**ware **L**ong-read **E**rror correction) and evaluated its performance on PacBio HiFi datasets from human and plant genomes. HALE matches the accuracy of state-of-the-art deep learning and heuristic-based methods while offering a more straightforward and theoretically grounded approach.

## 2 Preliminaries

Our proposed formulation (defined later in Section 3) is partly inspired by the *Hypercube 2-segmentation (H2S) problem* that was originally introduced by Kleinberg, Papadimitriou and Raghavan in 1998 for bi-clustering [5]. The problem statement is as follows.

### H2S Problem

Given a set of *n* vectors *x*_1_, *x*_2_, …, *x*_*n*_ in {0, 1}^*d*^, one needs to select two centers *c*_1_ and *c*_2_ in {0, 1}^*d*^ maximizing

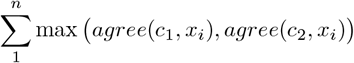

where function *agree*(*x, y*) counts on how many coordinates vectors *x* and *y* agree (which is also *d* minus the Hamming distance between *x* and *y*).

Another way to interpret the above problem is that we seek to partition *n* input vectors into two clusters such that the number of bit flips needed to make all the vectors agree within each cluster is minimum. H2S problem is known to be NP-hard [4].

A generalization of the above problem, where each vector can have one or more consecutive wildcard symbols at both ends, is commonly used in the context of reference-guided haplotype phasing [9, 15]. This generalized version of the problem is also referred to as the minimum error correction (MEC) problem. Solving the MEC problem is useful to determine the variants that are co-located on the same (maternal or paternal) haplotype. The MEC-based approach assumes the availability of a reference genome because each input vector is derived from an alignment of a read to the reference genome.

For *de novo* assembly, we cannot assume the availability of a reference genome. However, we assume the availability of a collection of overlaps between reads, which is a common first step in most long-read assemblers [12]. An overlap either implies (i) a sufficiently long approximate match between the suffix of a read and the prefix of another read or (ii) an approximate match of an entire read to a substring of another read. The overlaps identified using approximate string matching algorithms would comprise both true overlaps and false overlaps. The false overlaps arise from repetitive sequences within or across haplotypes. Accordingly, when considering a target read alongside its overlapping reads, we may have one, two, or even multiple “read clusters”, each corresponding to different haplotypes or genomic loci.

## 3 Problem Formulation

Our input is a multiple sequence alignment (MSA) between a substring of a target read and overlapping substrings from other reads (see Figure 1 for an illustration). We defer the details of how these MSAs are constructed to Section 5. Suppose that the MSA comprises *n* + 1 vectors in {*A, T, C, G*, −}^*d*^, where *n* is the number of overlapping substrings and *d* is the length of the MSA. We denote the vector in the MSA that represents the substring of the target read as *t*, and the set of all other vectors representing the overlapping substrings as *S*.

**Figure 1.**
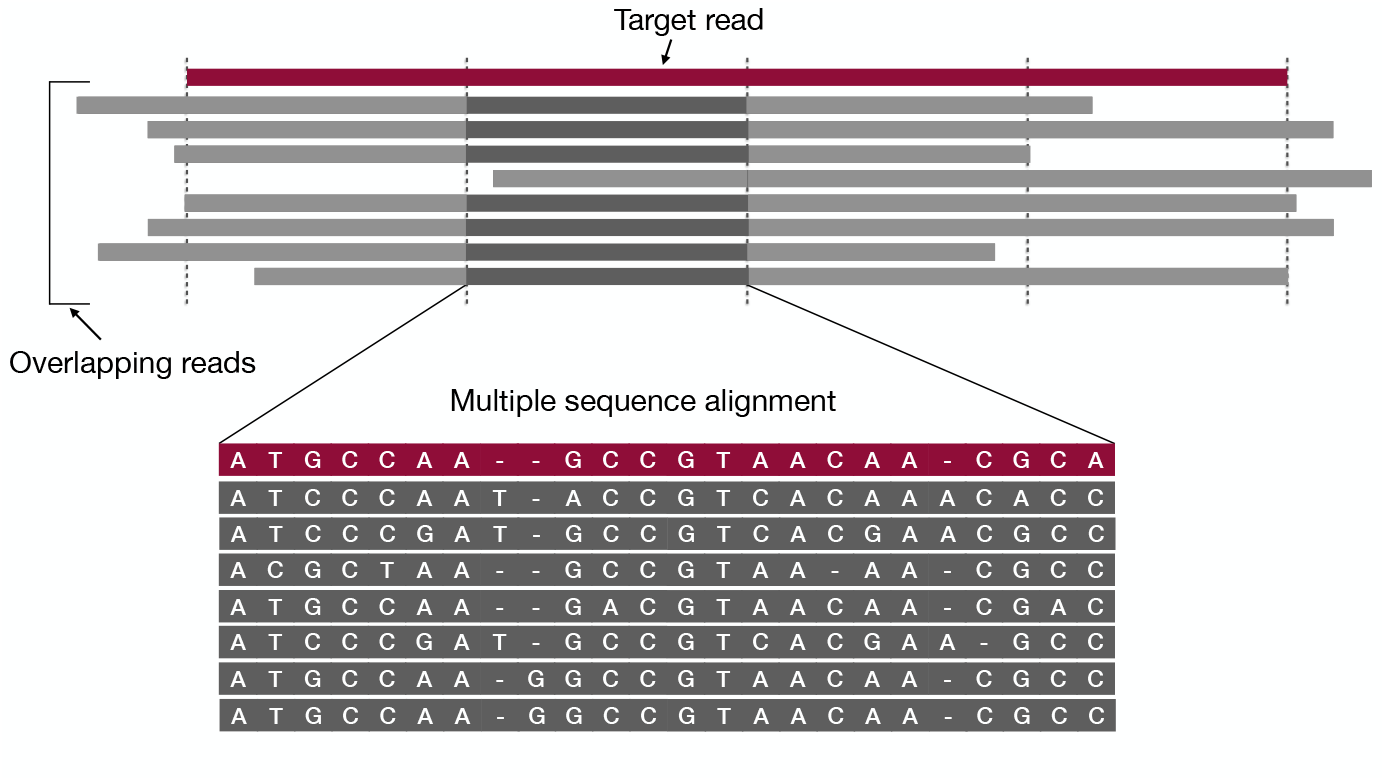
A toy example of the MSA input to our problem. MSA is computed between a substring of the target read and substrings of other overlapping reads. The target read is highlighted in red. Only those overlapping substrings are included in the MSA that have an end-to-end alignment with the substring of the target read (shown in dark grey). The above example has the target read split into four windows. In our implementation (discussed later), separate MSAs are generated for each window of the target read.

Some vectors in *S* represent true overlaps. They may differ from vector *t* due to sequencing errors in the reads. The remaining vectors differ from *t* due to both sequencing errors and haplotype-specific or repeat-specific variations. Accurately identifying the true overlaps is crucial for error correction, as it enables the use of a simple majority-vote strategy within the true overlaps. To achieve this, we introduce the following parsimony-based formulation aimed at selecting *k* vectors from *S* that, along with vector *t*, maximize a clustering score.

### Problem 1.

*Given a vector t in* {*A, T, C, G*, −}^*d*^, *a set S of n vectors in* {*A, T, C, G*, −}^*d*^, *and a positive integer k* ≤ *n, compute a subset S*′ ⊆ *S of cardinality k and a center c in* {*A, T, C, G*, −} ^*d*^ *such that* 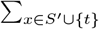 *agree* (*c, x*) *is maximum*.

We make a few remarks on this problem formulation. First, note that the symbol ‘−’ is not given any special treatment and is processed like any other character. Second, vector *c* in the solution satisfies *c*[*i*] = majority ({*x*[*i*] | *x* ∈ *S*′}) for all 1 ≤ *i* ≤ *d*, where ties are broken arbitrarily. Accordingly, *c* can be trivially computed once *S*′ is known. After solving Problem 1, vector *c* will be interpreted as the corrected version of vector *t*. Third, *k* is a user-defined parameter in Problem 1. Its value should depend on the sequencing coverage. One can approximately set *k* to the largest value such that every genomic interval of length *d* is sequenced at least *k* times with high probability.

Our problem formulation differs fundamentally from reference-guided haplotype phasing, e.g., WhatsHap [15]. Traditional reference-guided phasing aims to reconstruct two haplotypes by partitioning reads into two clusters. This clustering is disjoint and exhaustive, i.e., every read is assigned to exactly one of the two clusters. In contrast, we focus on identifying a subset of reads that corresponds to the haplotype and genomic region of the target read. The proposed formulation is more compatible with the *de novo* setting where the number of read clusters can vary.

## 4 Hardness Result

### Theorem 2.

*Problem 1 is NP-hard*.

Our proof technique is inspired by the technique used by Feige [4] to prove the hardness of the H2S problem. However, several arguments differ in our proof because we seek only one cluster in Problem 1 whereas the H2S problem seeks two clusters. Rather than proving the hardness of Problem 1 directly, we consider a related problem that slightly generalizes it.

### Problem 3.

*Given a set S of n vectors in* {−1, 1}^*d*^ *and a positive integer k* ≤ *n, compute a subset S*′ ⊆ *S of cardinality at least k that maximizes the ℓ*_1_ *norm of the vector sum, i*.*e*. 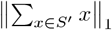.

In the Appendix (Lemma 7), we argue that Theorem 2 follows if Problem 3 is NP-hard. Our reduction to prove the hardness of Problem 3 uses Hadamard codes. A Hadamard code *H*_*M*_ of dimension *M* is a collection of *M* vectors in {−1, 1}^*M*^ such that every two vectors are orthogonal. There are well-known polynomial-time algorithms to construct *H*_*M*_ recursively when *M* is a power of two [19]. Therefore, we assume that *M* is a power of two. In the following lemma, we restate a useful property of Hadamard codes from [4].

### Lemma 4.

*Consider an arbitrary set of distinct vectors from the Hadamard code H*_*M*_. *The ℓ*_1_ *norm of their sum is at most M* ^3*/*2^.

### Lemma 5.

*Problem 3 is NP-hard*.

**Proof**. We reduce from the MAX-CUT problem. In the MAX-CUT problem, one is given an undirected graph, say *G*(*V, E*), and is asked to find a subset *V*_1_ of vertices that maximises the number of edges between *V*_1_ and its complement *V* \ *V*_1_.

We will convert *G* into an instance of Problem 3. Let us assign an arbitrary orientation to each edge of *G*. We will assume an integer parameter *M* (we will later find an appropriate value of *M* which is polynomial in the size of the graph).

We build a set *S* comprising *n* = *M* |*V* |vectors of dimension *d* = *M* |*E*|. For every vertex *v*_*i*_ of *G*, we introduce *M* vectors *v*_*i*,1_, …, *v*_*i,M*_ each of length *M* |*E*| (see Figure 2 for an illustration). Each vector is divided into |*E*| blocks of length *M* each. In each of the *n* vectors, in every block *B*_*e*_ associated with an edge *e*:

**Figure 2.**
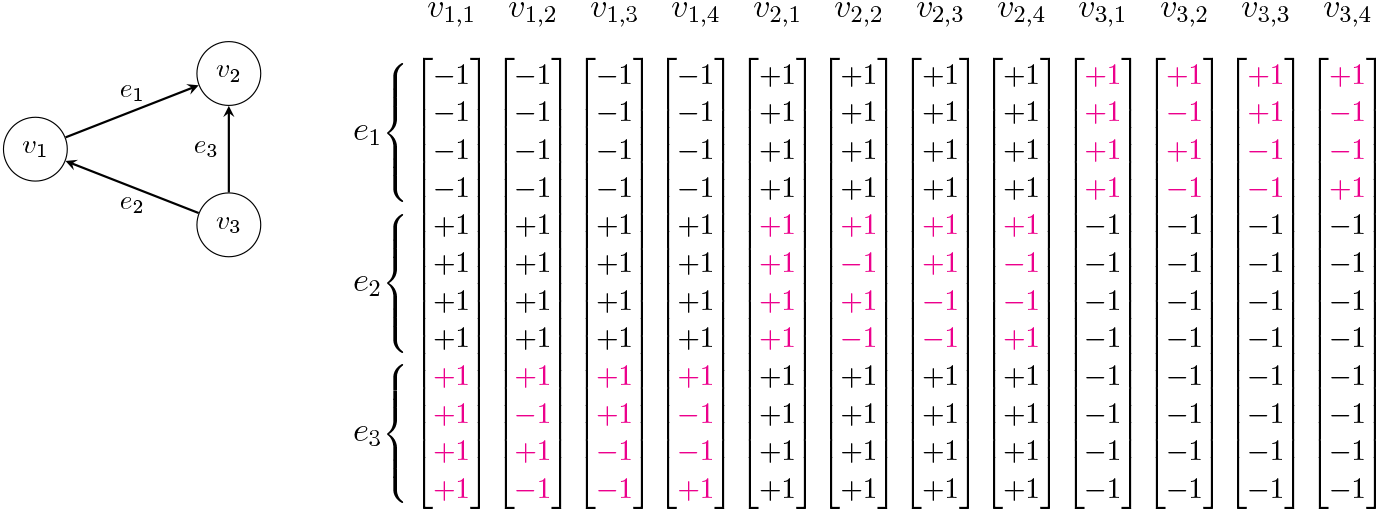
(Left) A directed graph. (Right) The resulting set of vectors when *M* = 4. The Hadamard codes are shown in pink. There are *M* |*V* | vectors, each of dimension *M* |*E*|.

1. If *v*_*i*_ is head of edge *e*, then all entries of *B*_*e*_ are +1.
2. If *v*_*i*_ is tail of edge *e*, then all entries of *B*_*e*_ are −1.
3. If *v*_*i*_ is not incident with edge *e*, then the entries of *B*_*e*_ in vector *v*_*i,j*_ (for 1 ≤ *j* ≤ *M*) are the *j*^*th*^ vector of Hadamard code *H*_*M*_.

Lastly, we set the parameter *k* = |*M*| (*V /*2).

**Yes** instances. Suppose that *G*(*V, E*) admits an optimal solution (*V*_1_, *V*_2_) having *c* edges between *V*_1_ and *V*_2_. WLOG, we can assume that |*V*_1_| ≥ |*V*_2_|, and hence |*V*_1_| ≥ |*V*| */*2. Consider one (possibly sub-optimal) solution to the instance of Problem 3 that selects all *M* |*V*_1_| vectors from *S* that correspond to vertices in *V*_1_. Let us denote the set of these *M* |*V*_1_| vectors as *X*.

A lower bound on the *ℓ*_1_ norm of the vector sum, i.e., 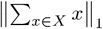, can be obtained as follows. Recall that each vector in *X* has |*E*| blocks. Consider a block associated with an edge *e* that is in the cut. There is exactly one endpoint of this edge in *X*, and all the *M* vectors corresponding to this endpoint (vertex) are monochromatic blocks of length *M*, which together contribute a total of *M* ^2^ to the *ℓ*_1_ norm. This can be partially offset by other blocks. But the block of each vertex not incident with *e* can offset at most *M* ^3*/*2^ of the *ℓ*_1_ norm (using Lemma 4). For an edge *e*, there are |*V*| − 2 vertices not incident with *e*. This makes the total contribution of the blocks corresponding to edge *e* to the *ℓ*_1_ norm at least *M* ^2^ (|*V*| 2)*M* ^3*/*2^. As there are *c* edges in the cut, the value of this solution is at least *c*(*M* ^2^ − (|*V*| − 2)*M* ^3*/*2^). (The value is actually higher because the blocks associated with the edges not in the cut also contribute to the *ℓ*_1_ norm, but we ignore this further tightening of the bound.)

**No** instances. Suppose that *G*(*V, E*) admits an optimal solution with *< c* edges in the cut. Consider an arbitrary subset of vectors from *S* with cardinality at least *k*. Call this subset of vectors *X*_1_. This selection corresponds to a *fractional* partition (*V*_1_, *V*_2_) of *G* where the extent to which vertex *v*_*i*_ is in partition *V*_1_ is equal to the fraction of its vectors in *X*_1_. Let *z*_*i*_ denote this extent for vector *v*_*i*_. Similarly, let 1 − *z*_*i*_ be the extent to which the vertex *v*_*i*_ is in partition *V*_2_. For an edge *e* = (*v*_*i*_, *v*_*j*_), the extent to which it is cut is *y*_*e*_ = |*z*_*i*_ − *z*_*j*_ |.

For an arbitrary edge *e*, the monochromatic blocks associated with its two endpoints together contribute *M* ^2^*y*_*e*_ to the *ℓ*_1_ norm of *X*_1_. The blocks associated with vertices that are not endpoints of *e*, each contribute at most *M* ^3*/*2^ towards the *ℓ*_1_ norm of *X*. Summing up over all edges and all blocks, the value of any solution is at most ∑_*e*_ (*M* ^2^*y* + (|*V*| − 2)*M* ^3*/*2^ = *M* ^2^ ∑_*e*_ *y*_*e*_ +(|*V*| − 2) |*E*| *M* ^3*/*2^. Further,∑ _*e*_ *y*_*e*_ ≤ *c* − 1. This is because it is possible to change a fractional cut into an integer cut which is at least as large (see Lemma 13 in Appendix). **Summary**. If we subtract the upper bound for *no* instances from the lower bound for *yes* instances, it follows that the *yes* instance leads to an higher value than the *no* instance if

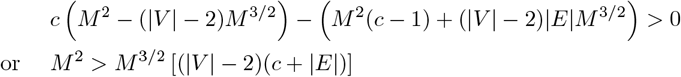

Taking *M >* 4|*E*|^2^|*V* |^2^ suffices. ◀

## 5 Algorithm and Implementation

This section outlines the details of our error-correction algorithm, HALE. Since computing exact MSA and solving Problem 1 are both NP-hard, we incorporated heuristics in our implementation. At the preprocessing stage, we use Minimap2 [7] to compute overlaps among all reads. Thus, for each read, we have a set of its pairwise alignments with other reads. We perform error correction of each read independently. In the following, we refer to the read being corrected as target read.

### Step 1: Constructing multiple sequence alignments

To construct MSAs, we adapted code from the Herro software [18]. We briefly summarize the approach here. Herro employs the well-known star alignment heuristic, where the target read is assumed to be the ‘center of the star’. The MSA is constructed progressively, i.e., adding each overlapping read to the MSA one by one using its precomputed pairwise alignment with the target read. For further details on the star alignment heuristic, see [17].

Once the full MSA is constructed, it is divided into contiguous, non-overlapping windows of fixed length *w* (Figure 1). Each window represents an MSA of a substring of the target read and its overlapping substrings. Henceforth, each windowed MSA is processed independently. Within each window, alignments from overlapping reads which do not fully span the window are ignored (Figure 1). We use the same window length *w* = 4096 as Herro.

If there are less than two alignments from overlapping reads in a window, which can happen occasionally in regions with low coverage, then Herro chops and removes the corresponding substring of the target read instead of leaving it uncorrected. We do the same in HALE. Note that this procedure may split the target read. The rationale is that the uncorrected portions of a read may negatively affect the quality of genome assembly, hence they are removed.

### Step 2: Sampling overlaps based on sequence identity

The count of overlapping substrings within a window is influenced by the sequencing coverage of the loci linked to the target read and how repetitive those regions are within the genome. To maintain computational efficiency of next steps, we retain only top-*n* overlapping substrings in each MSA. Preference is given to those overlapping substrings that have higher sequence identity^1^ with the target read. If fewer than *n* overlaps are available, we retain all. In HALE, the default value of parameter *n* is set to 20.

### Step 3: Sampling informative alignment columns

Following the approach of Herro [18] and Hifiasm [2], we identify *informative* alignment columns in the MSA. A column is considered informative if at least two distinct characters from the set {*A, T, C, G*,−}occur in that column with a minimum frequency of 3. See Figure 3 for an illustration. Only a small fraction of alignment columns are informative in practice. Informative columns typically indicate either biological variation among the aligned substrings or systematic sequencing errors, e.g., homopolymer indel errors. In contrast, non-informative columns are those where alignments largely agree. These columns have limited value for identifying true overlaps in Problem 1. Thus, we remove all non-informative columns from the MSA and use the modified MSA as input to Problem 1.

**Figure 3.**
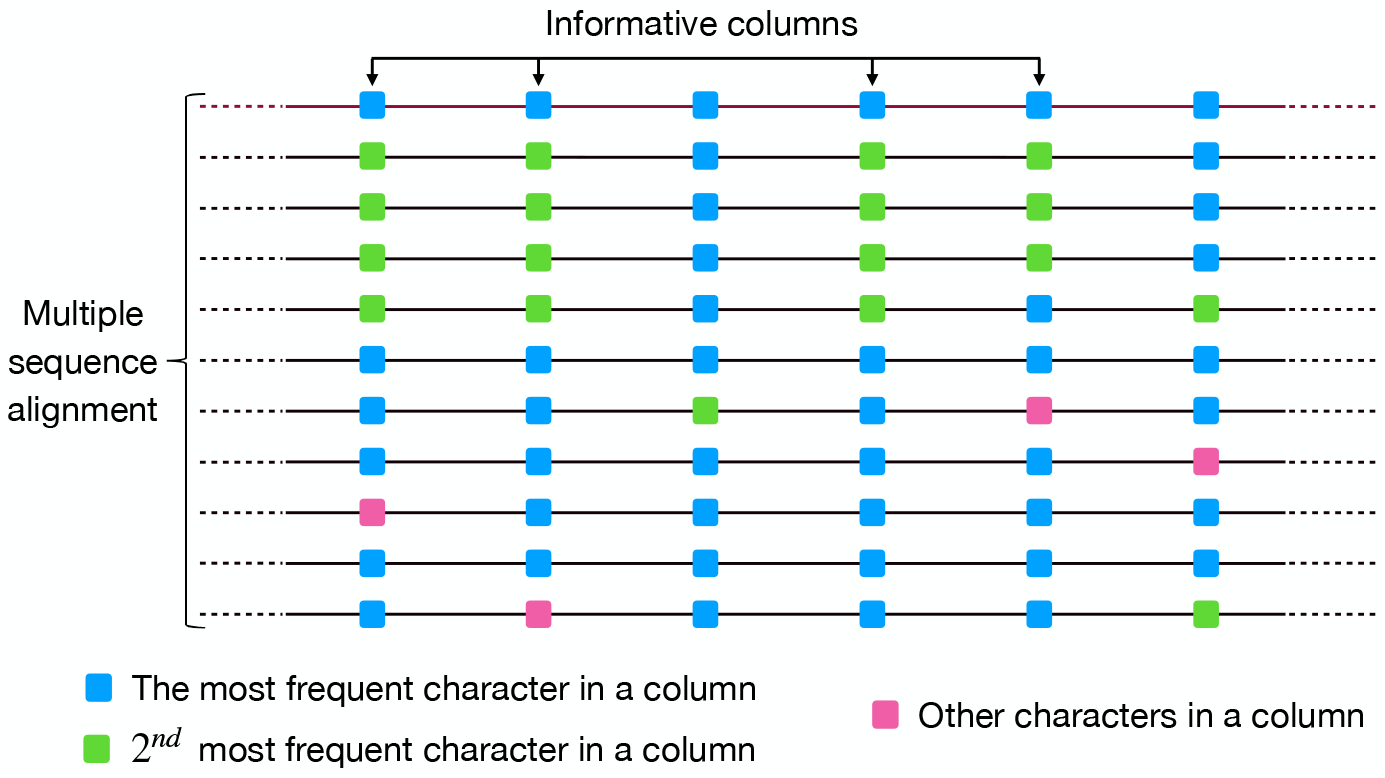
Informative columns in a multiple sequence alignment. In each column of the above MSA, blue boxes represent the most frequent character from {*A, T, C, G*, −} in that column. The green boxes represent the second most frequent character in a column. The pink boxes represent any other character that is neither the most nor the second most frequent. Based on the definition of informative columns, four out of six columns in the above example are informative.

### Step 4: Solving Problem 1

A polynomial-time algorithm is unlikely for Problem 1 because we have established its hardness in Section 4. We use a direct brute-force approach to find an exact solution. In the MSA input to Problem 1, we have a vector *t* in {*A, T, C, G*, −}^*d*^ (representing a subsequence of target read) along with *n* other vectors in {*A, T, C, G*, −}^*d*^ (from the overlapping reads). Here, *d* denotes the number of columns in a given MSA. We solve Problem 1 by exhaustively evaluating all 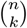 possible ways of selecting a subset of *k* vectors from the total of *n*. For a given selection of *k* vectors, an optimal value of the cluster center *c* at the *I* coordinate is the character that occurs most frequently at that position among the *k* vectors and the target vector *t*. Although the runtime of this algorithm is 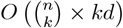, this brute-force solution is still practical due to the reduction in the size of MSA at Steps 2 and 3. There also exists a dynamic programming algorithm to solve Problem 1 (details omitted for brevity). We leave its implementation and evaluation for future work.

The default value of parameter *k* in HALE is set to *n/*3. This choice is based on our empirical observation that at least one-third of the overlapping substrings typically originate from the same haplotype as the target read (assuming ploidy ≤2). Choosing an appropriate value for *k* involves a trade-off: a significantly larger *k* can lead to the selection of reads with false overlaps, while a much smaller *k* leads to an insufficient number of reads to reliably compute a consensus.

We introduce an additional optimization. Note that Problem 1 treats all MSA columns with equal importance. To further improve the quality of results, we solve a weighted version of Problem 1 defined below. Suppose that weight function *W* assigns a non-negative weight *W* (*i*) to each column *i* ∈ {1, 2, …, *d*}and function *match*(*α, β*) returns 1 if characters *α* and *β* match, and 0 otherwise.

#### Problem 6.

*Given a vector t in* {*A, T, C, G*, −}^*d*^, *a set S of n vectors in* {*A, T, C, G*, −}^*d*^, *and a positive integer k* ≤ *n, compute a su bset S*′ ⊆ *S of cardinality k and a center c in* {*A, T, C, G*, −}^*d*^ *such that* 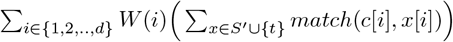 *is maximum*.

Our motivation to introduce the weights is as follows. A majority of errors in long reads are insertions and deletions, especially in homopolymers regions [11]. Accordingly, informative columns where the most frequent and the second most frequent characters are from {*A, T, C, G*} are likely to represent true biological variation. We assign a weight 1 to such columns. In contrast, other informative columns where either the most frequent or the second most frequent character is ‘−’ may correspond to either true indel variation or sequencing artifacts. We assign a weight 1*/*10 to these columns. The choice of this parameter was based on our empirical observations. Developing a more principled approach for determining the weight parameters is left for future work.

### Step 5: Correcting the target read

Recall that before sampling alignment columns in the MSA (Step 3), we had a substring of the target read aligned with overlapping substrings from other reads, forming an MSA of length *w*, where *w* is the window length parameter. The alignment of the substring from the target read corresponds to a vector in {*A, T, C, G*, −} ^*w*^. We update this vector as follows. For each non-informative column, where biological variation is not expected, we replace the character with the most frequent character in that column, which is equivalent to taking a majority vote. In all informative columns, we use the the optimal cluster center *c*, as determined by solving Problem 1, to update the characters. Finally, we generate the corrected version of the target read substring by removing any gap symbols (‘− ‘) from the updated vector.

## 6 Results

We evaluated the performance of HALE using publicly-available PacBio HiFi sequencing datasets. The commands and software versions to reproduce the results below are available in Appendix Tables 3 and 4. The HALE source code is accessible at https://github.com/at-cg/HALE.

### Datasets

We utilized two sequencing datasets, one from a diploid HG002 human sample^2^, and the other from a haploid Col-0 *Arabidopsis thaliana* sample^3^. Following the benchmarking approach used in Herro, we restricted our evaluation to reads that are at least 10 kbp in length. Reads shorter than 10 kbp were removed.

To reduce runtime of the experiments involving the human dataset, we sampled reads from chromosome 9 of the HG002 genome. Chromosome 9 contains a high abundance of satellite DNA sequences (long arrays of tandem repeats) which makes it one of the more challenging chromosomes to assemble [14]. To isolate reads from chromosome 9, we aligned all reads to HG002v1.1 assembly^4^ (GCA_018852605.3, GCA_018852615.3) and retained those reads whose primary alignments mapped to either the paternal or maternal haplotype of chromosome 9. To evaluate the impact of sequencing coverage on error correction performance, we created two versions of HG002 and *A. thaliana* datasets, one with 40× coverage and the other with 60× coverage by randomly sampling reads.

### Evaluated algorithms

We compared HALE against two state-of-the-art read correction tools, Herro [18] and Hifiasm [2]. Herro is a deep learning method that was primarily developed for correcting nanopore sequencing reads. Since a pretrained model for HiFi reads is not available in Herro, we used its existing model trained on Oxford Nanopore (R10) data. We assumed that the ONT-trained model generalizes to HiFi reads because HiFi reads are relatively more accurate and exhibit fewer systematic errors.

In addition to these baseline methods, we also compared HALE with two simplified correction methods. As a reminder, HALE performs a majority vote on non-informative alignment columns while using a more sophisticated technique based on solving Problem 1 in the informative alignment columns (Section 5). The naive methods employ the same heuristics described in Section 5, but bypass solving Problem 1 (Step 4).

In the first naive approach (Naive-1), a majority vote is performed across all alignment columns, including informative ones, during Step 5. In the second naive approach (Naive-2), the original values in the aligned target read are preserved at informative columns, while a majority vote is applied only to non-informative columns. This comparison allows us to isolate the benefit of solving Problem 1 as an optimization step in HALE.

### Evaluation metrics

For real sequencing datasets, there is no ground truth available on the errors made by the sequencing instrument. Thus, for both HG002 and *A. thaliana* datasets, we rely on high-quality references on which the reads can be mapped. For HG002 datasets, we use HG002v1.1 assembly (GCA_018852605.3, GCA_018852615.3) released by the Telomere-to-telomere consortium. For *A. thaliana* datasets, we use a high-quality assembly^5^ from [21].

To evaluate read correction performance, we aligned raw reads as well as reads corrected by different methods to their respective references using Minimap2. If a read aligns to multiple regions, then only its primary alignment was considered. We used BamConcordance [22] to examine the alignments. BamConcordance generates an empirical log-scaled measure of base error probabilities (Q-concordance) for each read. The formula used for calculating Q-concordance is −10 log_10_(1 − identity), where identity refers to the sequence identity observed in the read alignment. As an example, Q-concordance 20 corresponds to a sequence identity of 99%, indicating that 1% of the bases are erroneous. In addition to the Q-concordance statistics, we also report the percentage of reads containing at least *x* mismatch (similarly, *x* indel) errors, for *x* ∈ {1, 2, 3}, before and after correction.

As noted earlier in Section 5, the approach used in HALE and Herro discards a segment of a read if it cannot be reliably corrected due to insufficient overlapping read support. For a fair comparison, we evaluated only those reads that were fully corrected (i.e., not chopped) by all methods. Without this, HALE and Herro would have an unfair advantage over Hifiasm.

### Error correction performance

We show a comparison of the accuracy of raw reads and those corrected by HALE, Herro, and Hifiasm, respectively, in Figure 4a. We achieved comparable accuracy as Herro and Hifiasm. All three correction methods led to more than a 10-fold improvement in read accuracy relative to the raw, uncorrected reads.

**Figure 4.**
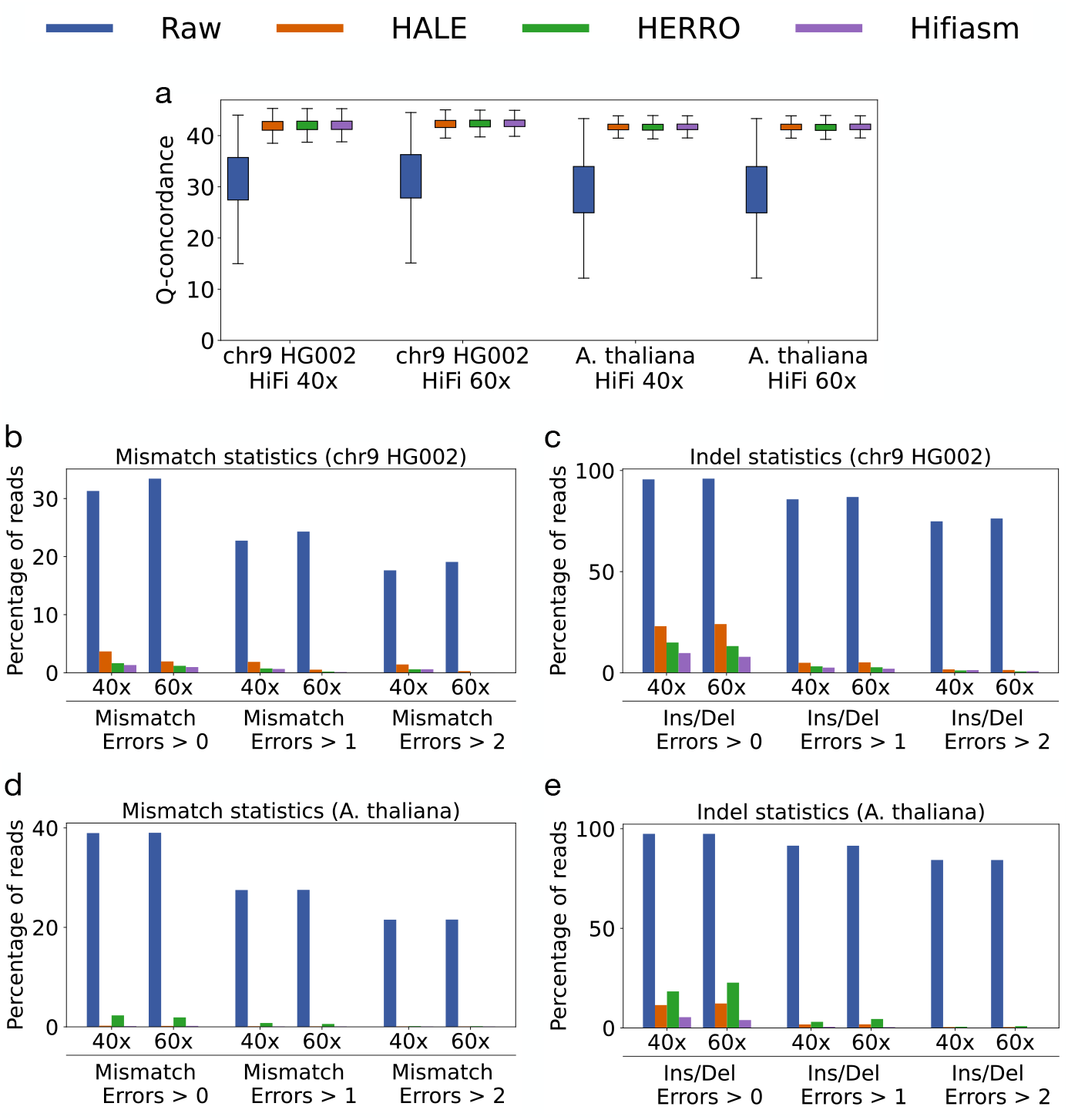
Accuracy of reads from chr9 HG002 and *A. thaliana* sequencing datasets at 40*×* and 60*×* coverage. The plots compare the accuracy of raw reads with those corrected by HALE, Herro, and Hifiasm. Table 1 in Appendix shows the exact values plotted in Figures 4b - 4e.

Next, we separately assessed mismatch and indel errors in the corrected reads. In Figure 4b, we highlight that the fraction of HG002 chr9 reads with one or more mismatch errors significantly drops after error correction. This result suggests that all three methods are effective in preserving biological variations in the reads. At 40 × sequencing depth, HALE exhibits a marginally higher percentage of reads containing at least *x* mismatch errors (*x* ∈ 1, 2, 3) compared to Herro and Hifiasm. However, this difference diminishes at 60× depth. We also tested HALE using reads from HG002 chromosomes 13 and 18, observing a consistent trend (Appendix Figure 6).

**Table 1.** The exact values plotted in Figures 4b - 4e.

| Dataset | Error type | # Errors | Percentage of reads |  |  |  |
| --- | --- | --- | --- | --- | --- | --- |
|  |  |  | Raw | HALE | Herro | Hifiasm |
| HG002 chr9 40x | Mismatch | >0 | 31.3 | 3.7 | 1.6 | 1.3 |
|  |  | >1 | 22.7 | 1.9 | 0.7 | 0.6 |
|  |  | >2 | 17.6 | 1.4 | 0.6 | 0.6 |
|  | Indel | >0 | 95.6 | 22.6 | 14.9 | 9.7 |
|  |  | >1 | 85.8 | 4.9 | 3.1 | 2.5 |
|  |  | >2 | 74.9 | 1.6 | 1.1 | 1.3 |
| HG002 chr9 60x | Mismatch | >0 | 33.4 | 1.9 | 1.2 | 1.0 |
|  |  | >1 | 24.3 | 0.5 | 0.2 | 0.1 |
|  |  | >2 | 19.1 | 0.3 | 0.0 | 0.0 |
|  | Indel | >0 | 96.0 | 23.1 | 13.2 | 7.9 |
|  |  | >1 | 86.9 | 5.1 | 2.7 | 2.0 |
|  |  | >2 | 76.3 | 1.3 | 0.6 | 0.7 |
| *A. thaliana* 40x | Mismatch | >0 | 39.0 | 0.2 | 2.3 | 0.1 |
|  |  | >1 | 27.5 | 0.1 | 0.8 | 0.0 |
|  |  | >2 | 21.5 | 0.1 | 0.1 | 0.0 |
|  | Indel | >0 | 97.4 | 11.5 | 18.3 | 5.4 |
|  |  | >1 | 91.5 | 1.7 | 3.0 | 0.5 |
|  |  | >2 | 84.3 | 0.4 | 0.6 | 0.1 |
| *A. thaliana* 60x | Mismatch | >0 | 39.0 | 0.2 | 1.9 | 0.2 |
|  |  | >1 | 27.5 | 0.1 | 0.6 | 0.1 |
|  |  | >2 | 21.6 | 0.1 | 0.1 | 0.0 |
|  | Indel | >0 | 97.4 | 12.3 | 22.7 | 3.9 |
|  |  | >1 | 91.5 | 1.8 | 4.5 | 0.4 |
|  |  | >2 | 84.3 | 0.4 | 0.9 | 0.1 |

In the case of *A. thaliana* dataset, which is a haploid genome sample, HALE is again comparable to the best performing method (Figures 4d, 4e). Herro lags slightly behind HALE and Hifiasm here, possibly because its model was trained using a diploid sample and nanopore reads [18].

We also manually checked and visualized a few reads in IGV [16] where HALE failed to correct one or more errors. In some cases, we found that uneven sequencing coverage in some genomic regions resulted in MSA windows where fewer than *k* true overlaps were present. As a result, our algorithm, constrained by the fixed cutoff of *k*, selected substrings from false overlaps for correction.

### Comparison with naive algorithms

Solving Problem 1 (Section 3) is an important component of HALE as it enables us to preserve the haplotype-specific variation in reads. We compared HALE with two simplified versions of our algorithm, Naive-1 and Naive-2, respectively, which omit this optimization step.

#### Comparison with Naive-1

On both HG002 and *A. thaliana* datasets, the Naive-1 algorithm achieves Q-concordance scores comparable to HALE (Figure 5a). These results, especially on HG002 diploid datasets, surprised us initially. Recall that the Naive-1 algorithm applies the majority vote on all alignment columns, irrespective of whether they are informative or not. We further assessed the mismatch and indel error statistics. We show that HALE provides a significant improvement over the Naive-1 algorithm in correcting mismatch errors (Figure 5b). This confirms that a direct consensus approach eliminates haplotype variation, whereas our optimization based on solving Problem 1 helps to preserve them.

**Figure 5.**
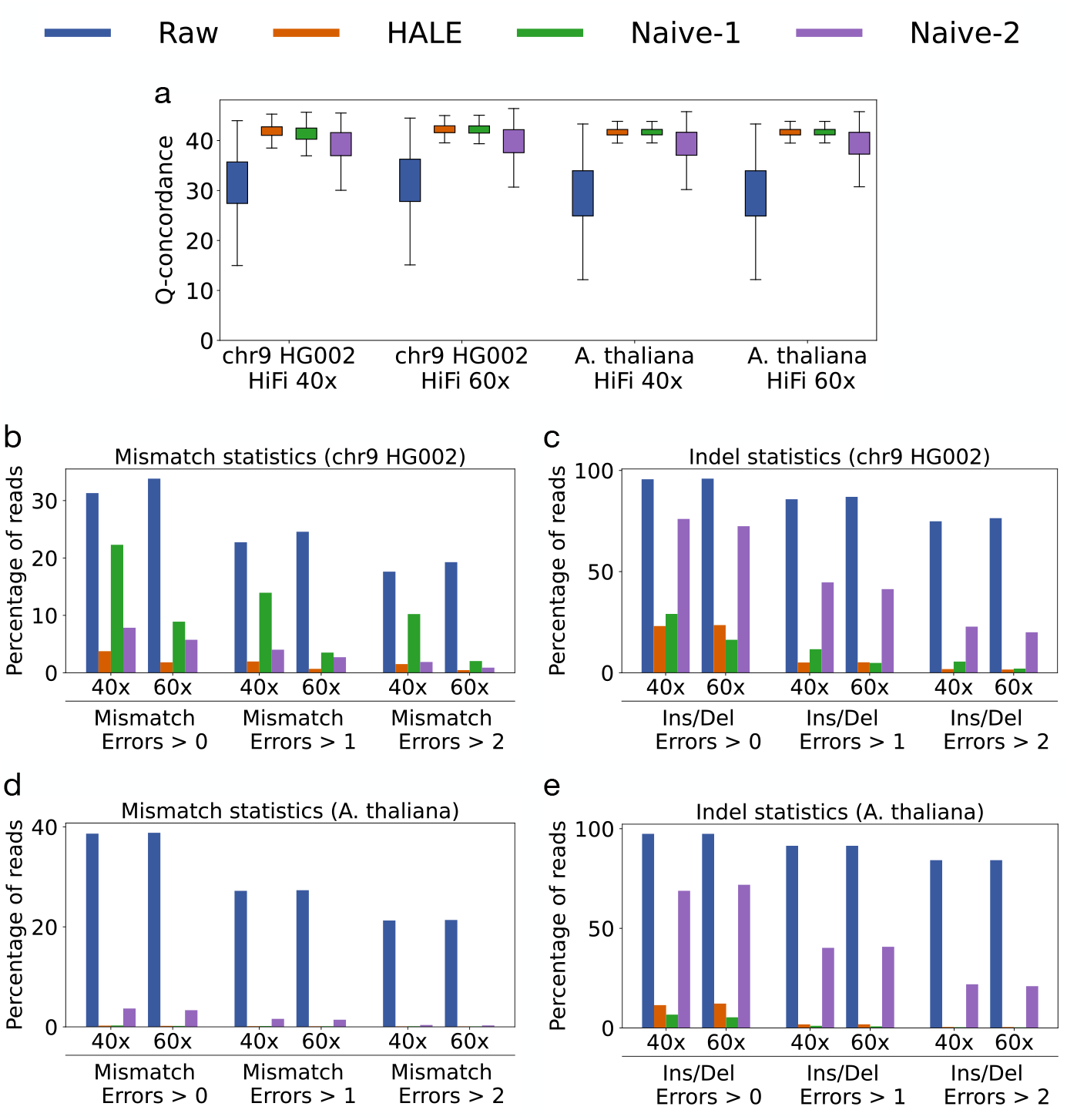
Accuracy of reads from chr9 HG002 and *A. thaliana* sequencing datasets at 40*×* and 60*×* coverage. The plots compare the accuracy of raw reads with those corrected by HALE, Naive-1, and Naive-2 algorithms. Table 2 in Appendix shows the exact values plotted in Figures 5b - 5e.

Despite this significant advantage in mismatch statistics, the Q-concordance scores in Figure 5a do not reflect a proportionate gain. This is because indel errors are the dominant error type in HiFi reads. As shown in Figure 5c, HALE performs comparably to Naive-1 when it comes to correcting indel errors.

**Table 2.** The exact values plotted in Figures 5b - 5e.

| Dataset | Error type | # Errors | Percentage of reads |  |  |  |
| --- | --- | --- | --- | --- | --- | --- |
|  |  |  | Raw | HALE | Naive-1 | Naive-2 |
| HG002 chr9 40x | Mismatch | >0 | 31.3 | 3.8 | 22.3 | 7.8 |
|  |  | >1 | 22.7 | 2.0 | 13.9 | 4.0 |
|  |  | >2 | 17.6 | 1.5 | 10.2 | 1.9 |
|  | Indel | >0 | 95.6 | 23.1 | 29.0 | 76.0 |
|  |  | >1 | 85.8 | 5.0 | 11.6 | 44.7 |
|  |  | >2 | 74.9 | 1.8 | 5.5 | 22.8 |
| HG002 chr9 60x | Mismatch | >0 | 33.4 | 1.8 | 8.9 | 5.7 |
|  |  | >1 | 24.3 | 0.7 | 3.5 | 2.7 |
|  |  | >2 | 19.1 | 0.4 | 2.0 | 0.9 |
|  | Indel | >0 | 96.0 | 23.5 | 16.3 | 72.4 |
|  |  | >1 | 86.9 | 5.1 | 4.8 | 41.3 |
|  |  | >2 | 76.3 | 1.6 | 2.0 | 20.0 |
| *A. thaliana* 40x | Mismatch | >0 | 39.0 | 0.2 | 0.3 | 3.7 |
|  |  | >1 | 27.5 | 0.1 | 0.1 | 1.6 |
|  |  | >2 | 21.5 | 0.1 | 0.1 | 0.4 |
|  | Indel | >0 | 97.4 | 11.4 | 6.7 | 68.8 |
|  |  | >1 | 91.4 | 1.8 | 1.0 | 40.2 |
|  |  | >2 | 84.3 | 0.5 | 0.4 | 21.9 |
| *A. thaliana* 60x | Mismatch | >0 | 39.0 | 0.2 | 0.2 | 3.3 |
|  |  | >1 | 27.5 | 0.1 | 0.1 | 1.4 |
|  |  | >2 | 21.6 | 0.1 | 0.1 | 0.3 |
|  | Indel | >0 | 97.4 | 12.2 | 5.3 | 71.8 |
|  |  | >1 | 91.4 | 1.8 | 0.8 | 40.7 |
|  |  | >2 | 84.3 | 0.4 | 0.3 | 21.0 |

Using the haploid *A. thaliana* dataset, the results are as expected. Naive-1 algorithm performs on par with HALE in terms of correcting mismatch errors (Figure 5d), and even slightly surpasses HALE when it comes to correcting indel errors (5e). A direct consensus-based approach works well on haploid datasets.

#### Comparison with Naive-2

We see a drop in the Q-concordance scores using the Naive-2 algorithm in both HG002 and *A. thaliana* datasets. This is because the Naive-2 algorithm makes no modification in a target read on informative alignment columns. In practice, a small fraction of informative alignment columns are classified as informative due to multiple sequencing errors occurring at the same coordinate of the target read and its overlapping reads. In such a case, Naive-2 will leave the erroneous base of the target read uncorrected. As a result, we see a much larger number of indel errors in Naive-2 (Figures 5c, 5e).

### Runtime comparison

Runtime comparisons between HALE, Herro, and Hifiasm are challenging for two main reasons. First, Hifiasm uses its own custom all-vs-all read overlapping algorithm that is tightly integrated inside its code, whereas HALE and Herro use an external tool, Minimap2 [7]. Second, Herro leverages GPU acceleration, whereas both HALE and Hifiasm run on CPUs. In the following, we report the wall-clock runtimes of all-vs-all read overlapping and error correction steps for HALE and Herro. For Hifiasm, we report the total end-to-end assembly time. We report the results using chr9 HG002 60× dataset.

Minimap2 used 26 minutes to compute all-vs-all overlaps using 64 CPU threads. HALE used 9 minutes for error correction using the same number of threads. In comparison, Herro required 10 minutes on a server with four NVIDIA A100 GPUs. Hifiasm completed the full assembly in 16 minutes using 64 CPU threads, with approximately 90% of that time spent on read overlapping and error correction steps. Based on these results, we conclude that HALE is computationally more efficient than Herro. Hifiasm is the fastest overall due to its highly optimized read overlapping implementation. As all-vs-all overlap computation is a runtime bottleneck while using HALE, using a faster read overlapper would reduce the runtime.

## 7 Discussion

Genome assembly tools often require a significant software engineering effort and rely heavily on heuristics that are often undocumented. It is important to investigate alternative algorithms that are grounded in solid theoretical principles. Error correction is one of the critical steps in genome assembly, especially in the context of diploid genomes, polyploid genomes, and metagenomes.

In this work, we proposed the first rigorous formulation for haplotype-aware error correction of long reads. The formulation is partly inspired from the bi-clustering formulation introduced in [5]. An extension of the same optimization framework is also used in reference-guided haplotype phasing tools such as WhatsHap [15]. Our proposed formulation is designed to identify true overlaps of a read from a given set of overlaps. Considering that the problem is NP-hard, we also discussed practical heuristics to make our algorithm scalable. We showcased the effectiveness of our algorithm using publicly available long-read datasets from a haploid *A. thaliana* genome and a diploid human genome. We showed that HALE achieves accuracy on par with state-of-the-art tools, including Herro (a deep learning-based method) and Hifiasm (which relies on intricate heuristics). Lastly, we isolated the benefit of using Problem 1 (Section 3) by comparing HALE with the other simplified versions of our algorithm.

A number of improvements are possible to the proposed optimization framework as well as our implementation. One limitation of our formulation is that we independently process windows of an MSA (Figure 1) rather than processing the entire MSA at once. This can be problematic when a window is particularly noisy or lacks sufficient haplotype-specific variation, making it difficult to select the correct subset of overlapping reads. Future work could explore alternative formulations that exploit the full length of long reads. Another limitation is our reliance on parameters such as *k* (count of overlapping reads to be selected) and the MSA window length. In this work, we primarily relied on our empirical observations to set these parameters. Developing either parameter-free approaches or a principled way to select these parameters which generalizes across haploid, diploid, and polyploid genomes would be an important next step.

Regarding the implementation of heuristics in HALE, the use of superior all-vs-all read aligners and MSA heuristics may further enhance read accuracy. Furthermore, it may be possible to improve the selection of informative alignment columns in an MSA such that a larger fraction of informative columns correspond to biological variation rather than sequencing errors. For example, indel variation in homopolymer regions is more likely to be a sequencing error than a biological variation [3].

HALE is theoretically promising, it is not yet a practical replacement for existing state-of-the-art tools. In future, we hope to continue improving HALE. We also hope to extend its applicability to nanopore sequencing reads, which have higher error rates than HiFi reads.

## Acknowledgements

The authors thank Bikram Kumar Panda, Sudhanva Shyam Kamath, Mile Sikic, and Paul Medvedev for providing useful feedback during the project.

## Appendix

### 1 Hardness Result: Supplementary

#### Lemma 7.

*The NP-hardness of Problem 3 implies NP-hardness of Problem 1*

**Proof**. Our proof uses a chain of arguments. We begin by defining Problem 8, a simplified version of Problem 1 where we have a single term in the objective function and a binary alphabet. Next, we introduce Problem 9, a modified version of Problem 3. To prove the lemma, we first prove Claim 10 to show that Problem 9 is NP-hard, then Claim 11 to establish polynomial-time equivalence between Problem 9 and Problem 8. Finally we complete the proof by proving Claim 12.

#### Problem 8.

*Given a set S of n vectors in* {0, 1} ^*d*^ *and a positive integer k* ≤ *n, compute a subset S*′ ⊆ *S of cardinality k and a center c in*{0, 1} ^*d*^ *such that* 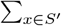 *agree*(*c, x*) *is maximum*.

#### Problem 9.

*Given a set S of n vectors in* {−1, 1} ^*d*^ *and a positive integer k* ≤ *n, compute a subset S*′ ⊆ *S of cardinality k that maximizes the ℓ*_1_ *norm of the vector sum, i*.*e*. 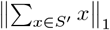.

#### Claim 10.

The NP-hardness of Problem 3 (Lemma 5) implies the NP-hardness of Problem 9.

**Proof**. Problem 9 differs slightly from Problem 3 because we seek a subset of size exactly *k* in Problem 9, whereas we seek a subset of size at least *k* in Problem 3. We prove Claim 10 using a simple polynomial-time reduction from Problem 3 to Problem 9. Given an instance of Problem 3 with *S* = {*x*_1_, …, *x*_*n*_} and parameter *k* = *α*, we construct *n* − *α* + 1 instances for Problem 9 using the same set of vectors *S* and parameter *k* = *α* + *i*, for all 0 ≤ *i* ≤ *n* − *α*. The maximum of the solution over all instances of Problem 9 is the solution to Problem 3. ◀

#### Claim 11.

Problems 8 and 9 are polynomial-time equivalent problems.

**Proof**. We consider a solution set of vectors *S*′ ⊆ {0, 1} ^*d*^ for Problem 8. Observe that the vector *c* must satisfy *c*[*i*] = majority ({*x*[*i*] | *x ∈ S*′}), where ties are broken arbitrarily. Let *ϕ* : {0, 1} → {−1, 1} be the coordinate-wise transformation defined by *ϕ*(0) = 1 and *ϕ*(1) = −1. For any vector *x* ∈ {0, 1} ^*d*^, let 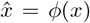 denote its image in {−1, 1}^*d*^. Let *Ŝ*′ ⊆ {−1, 1} ^*d*^ be the set of vectors *S*′ with *ϕ* applied to each vector. The objective function of Problem 8 can be written as

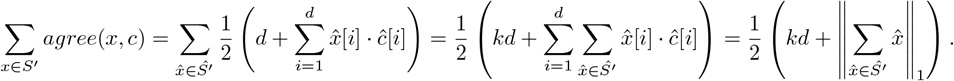

Thus, maximizing the objective function of Problem 8 is equivalent to maximising 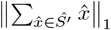.

Given a solution to Problem 8, i.e., a vector *c* ∈ {0, 1} ^*d*^ and subset of vectors *S*′ ⊆ {0, 1} ^*d*^ that maximizes Σ_*x*∈*S*′_ *agree*(*x, c*), we obtain a solution for Problem 9 by applying *ϕ*(·) component-wise to every vector in *S*′, creating the set *Ŝ* ⊆ {−1 1} ^*d*^. By our earlier argument, 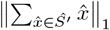 is maximised. Conversely, given a solution for Problem 9 of the form *Ŝ*′ ⊆ {−1, 1}^*d*^, we apply *ϕ*^−1^ to every vector in *Ŝ*′ to obtain a solution *S*′ ⊆ {0, 1}^*d*^ for Problem 8. The center vector *c* is obtained by taking a majority over each coordinate, and the objective function of Problem 8 (sum of agreement with *c*) is maximized. ◀

#### Claim 12.

The NP-hardness of Problem 8 implies the NP-hardness of Problem 1.

**Proof**. We prove this by polynomial-time reduction from Problem 8 to Problem 1 defined on alphabet Σ = {0, 1} (since this can be bijectively mapped to {*A, T*}). Given an instance of Problem 8 with set *S* = {*x*_1_, …, *x*_*n*_} and parameter *k* = *α*, construct *n* instances of Problem 1 where the *i*^*th*^ instance (*t*^*i*^, *S*^*i*^, *k* = *α* − 1) is constructed by considering *x*_*i*_ as the target vector *t*^*i*^, set *S*^*i*^ = *S* \ *x*_*i*_, and parameter *k* = *α* − 1. The maximum value solution of all *n* instances of Problem 1 is the solution to Problem 8. ◀

◀

#### Lemma 13.

*Given a fractional solution to the Max-Cut problem, there exists a rounding procedure that transforms it into an integral solution which is at least as large as that of the fractional solution*.

**Proof**. Let *z*_*i*_ ∈ [0, 1] be the given fractional assignment of vertex *v*_*i*_, for *i* ∈ [1, |*V* |]. The extent to which an edge (*v, v*) is cut is given by |*z*_*i*_ − *z*_*j*_|. Accordingly, the cut size of the given fractional partitioning is given by ∑ _(*i,j*) ∈ *E*_ |*z*_*i*_ − *z*_*j*_ |. Next, consider the following procedure to convert fractional assignment of vertices into an integral assignment. Let *t* be a threshold chosen uniformly at random from the interval [0, 1]. Assign vertex *v*_*i*_ to partition *V*_1_ if *z*_*i*_ *> t* and assign it to partition *V*_2_ otherwise. For any edge (*v*_*i*_, *v*_*j*_) ∈ *E*, the probability that it is cut is given by |*z*_*i*_ − *z*_*j*_| because this occurs precisely when *z*_*i*_ ≤ *t < z*_*j*_. Hence, the expected value of the cut size of our integral solution is ∑_(*v, v*) *E*_ |*z*_*i*_ *z*_*j*_|, which is equal to the cut size of the fractional cut. This expectation guarantees the existence of at least one threshold *t*^*^ [0, 1] for which the corresponding integral solution has value at least as large as the given fractional solution. Finally, observe that it suffices to check at only |*V*| + 2 candidate values for *t*: namely, at each *z*_*i*_ ∈ {*z*_1_, *z*_2_, …, *z* _|*V*|_}, as well as the endpoints 0 and 1. This is because the objective remains constant between any two consecutive values in the sorted sequence *z*_1_ *< z*_2_ *<* … *< z*_|*V* |_. Therefore, an optimal value *t*^*^ can be found in polynomial time by checking the solution at only these |*V* | + 2 points.◀

*Remark:* Lemma 13 was stated directly without a proof in [4].

### 2 Additional Results

**Figure 6.**
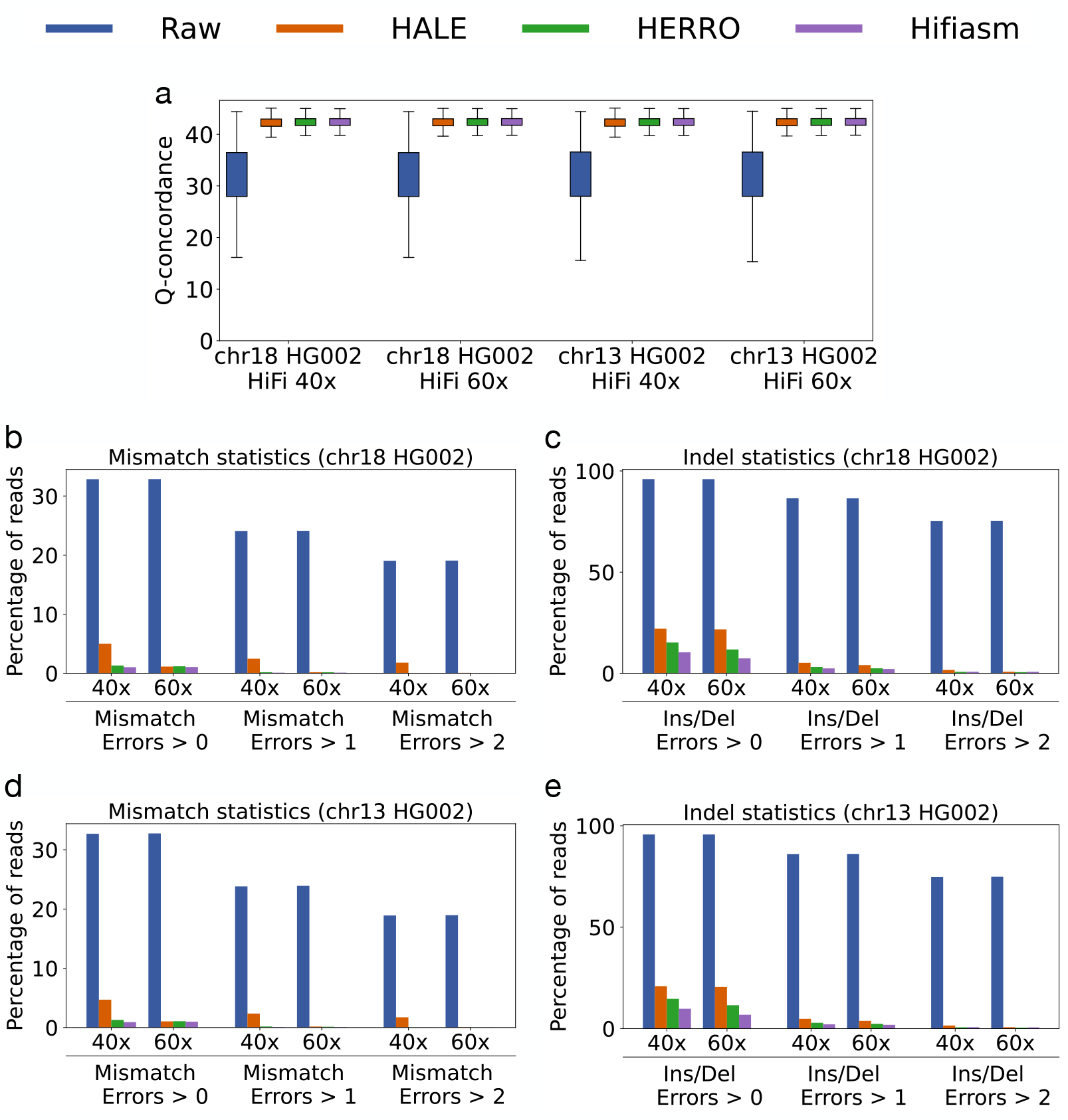
Accuracy of sequencing reads from HG002 chromosomes 18 and 13, at 40*×* and 60*×* coverage. The plots compare the accuracy of raw reads with those corrected by HALE, Herro, and Hifiasm.

### 3 Software Versions and Commands

**Table 3.**
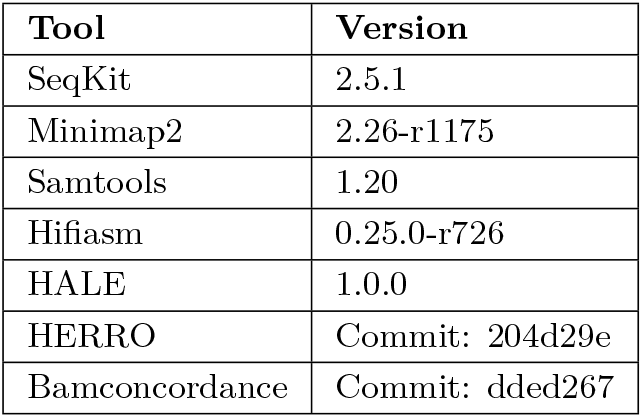
Versions of software used in our experiments.

| Tool | Version |
| --- | --- |
| SeqKit | 2.5.1 |
| Minimap2 | 2.26-r1175 |
| Samtools | 1.20 |
| Hifiasm | 0.25.0-r726 |
| HALE | 1.0.0 |
| HERRO | Commit: 204d29e |
| Bamconcordance | Commit: dded267 |

**Table 4.**
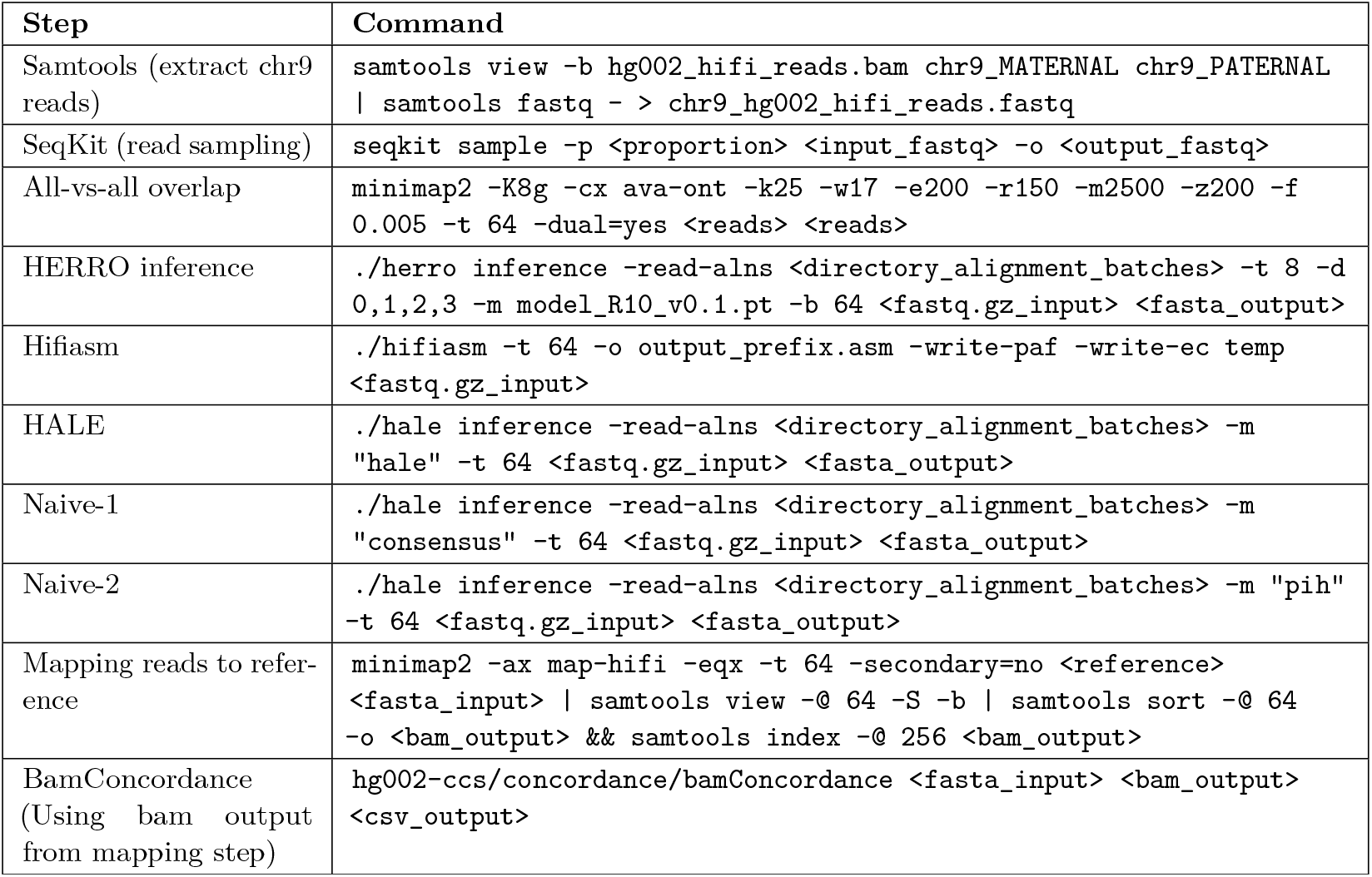
Commands used to run various tools.

## Footnotes

1 Given a pairwise alignment of two sequences, sequence identity is defined as the number of matching bases over the number of alignment columns.

2 https://s3-us-west-2.amazonaws.com/human-pangenomics/T2T/scratch/HG002/sequencing/hifirevio/

3 https://ngdc.cncb.ac.cn/gsa/browse/CRA004538/CRR302668

4 https://s3-us-west-2.amazonaws.com/human-pangenomics/T2T/HG002/assemblies/hg002v1.1.fasta.gz

5 https://ngdc.cncb.ac.cn/gwh/Assembly/21820/show

